# Optimal construction of a functional interaction network from pooled library CRISPR fitness screens

**DOI:** 10.1101/2022.08.03.502694

**Authors:** Veronica Gheorghe, Traver Hart

## Abstract

Functional interaction networks, where edges connect genes likely to operate in the same biological process or pathway, can be inferred from CRISPR knockout screens in cancer cell lines. Genes with similar knockout fitness profiles across a sufficiently diverse set of cell line screens are likely to be co-functional, and these “coessentiality” networks are increasingly powerful predictors of gene function and biological modularity. While several such networks have been published, most use different algorithms for each step of the network construction process. In this study, we identify an optimal measure of functional interaction and test all combinations of options at each step – essentiality scoring, sample variance and covariance normalization, and similarity measurement – to identify best practices for generating a functional interaction network from CRISPR knockout data. We show that Bayes Factor and Ceres scores give the best results, that Ceres outperforms the newer Chronos scoring scheme, and that covariance normalization is a critical step in network construction. We further show that Pearson correlation, mathematically identical to ordinary least squares after covariance normalization, can be extended by using partial correlation to detect and amplify signals from “moonlighting” proteins which show context-dependent interaction with different partners.

## Introduction

Functional interaction networks connect genes which operate in the same biological process or pathway. Systematic genetic interaction surveys in yeast showed that genes with similar genetic interaction profiles across dozens (Tong et al., 2001) to thousands (Costanzo et al., 2010; Costanzo et al., 2016) of strains showed high likelihood of functional interaction. In human cells, genome-scale pooled library knockdown (Hart et al., 2017) or CRISPR-mediated gene knockout screens (Wang et al., 2015) (Rauscher et al., 2018) (Boyle et al., 2018) (Kim et al., 2019) (Wainberg et al., 2021) enabled the comparison of gene loss of function fitness vectors across cell lines, which show the same tendency toward co-functionality. As the number of cell line CRISPR screens has grown (Broad Institute, 2019), these coessentiality networks have become a powerful method for inferring gene function and understanding the modular architecture of the cell (Kim et al., 2022a).

Though informative, there has to date been no systematic evaluation of the quality of each of these networks – nor even consensus on how to measure quality. There are numerous algorithms for transforming raw gRNA read count data from pooled library CRISPR screens into quantitative measurements of gene fitness effect, including Bagel2 (Kim and Hart, 2021), Castle (Morgens et al., 2017), Ceres (Meyers et al., 2017), Chronos (Dempster et al., 2021), JACKS (Allen et al., 2019), MAGECK (Li et al., 2014), and a Z-score approach (Lenoir et al., 2021) optimized for finding genes with positive instead of negative knockout fitness effects.

After combining gRNA-level fold changes into gene-level scores, each published approach uses different sample-level variance normalization. Kim et al (Kim et al., 2019) use quantile normalization with no reference distribution, which resets the value of each gene to the mean of all gene fitness scores at that rank. Boyle et al (Boyle et al., 2018) subtract the top principle components of the fitness matrix of a set of reference nonessential genes (olfactory receptors), to remove artifactual components. Wainberg et al (Wainberg et al., 2021) go further and implement a covariance whitening method based on Cholesky decomposition, which normalizes both variance and covariance and minimizes the effect of uneven sample distribution – a potentially serious source of bias, since, for example, there are more than ten times as many lung cancer cell lines in DepMap as prostate cancer lines. The similarity of gene vectors is frequently measured by Pearson correlation coefficient (PCC), though Wainberg et al argue that inflated P-values from PCC lead to numerous false positives and that ordinary least squares (OLS) after Cholesky whitening – collectively Generalized Least Squares (GLS) – is a better approach.

In this study, we systematically compare combinations of essentiality scoring algorithms, variance and covariance normalization methods, and similarity measures to determine an optimal strategy for building a functional interaction network from coessentiality data. We show that results are highly dependent on data processing steps, and that covariance whitening is a critical step in improving the predictive power of networks. We further show that, after covariance normalization, PCC and OLS are mathematically identical, and demonstrate how PCC and partial correlation can be employed to extract context-dependent interactions from global coessentiality networks.

## Results & Discussion

To assess an optimal functional interaction network from coessentiality data, we first acquired gene knockout fitness data from CRISPR knockout screens from the Cancer Dependency Map project. A typical network construction pipeline involves converting raw read count data into a matrix of gene knockout fitness scores, performing variance and/or covariance normalization across samples, and measuring similarity of all pairs of gene fitness profiles across samples (Figure 1). We constructed networks using all combinations of alternatives at each step and evaluated the quality of each network using a log likelihood framework that measures the enrichment for gene pairs that belong to the same annotated biological process or pathway.

**Figure 1.**
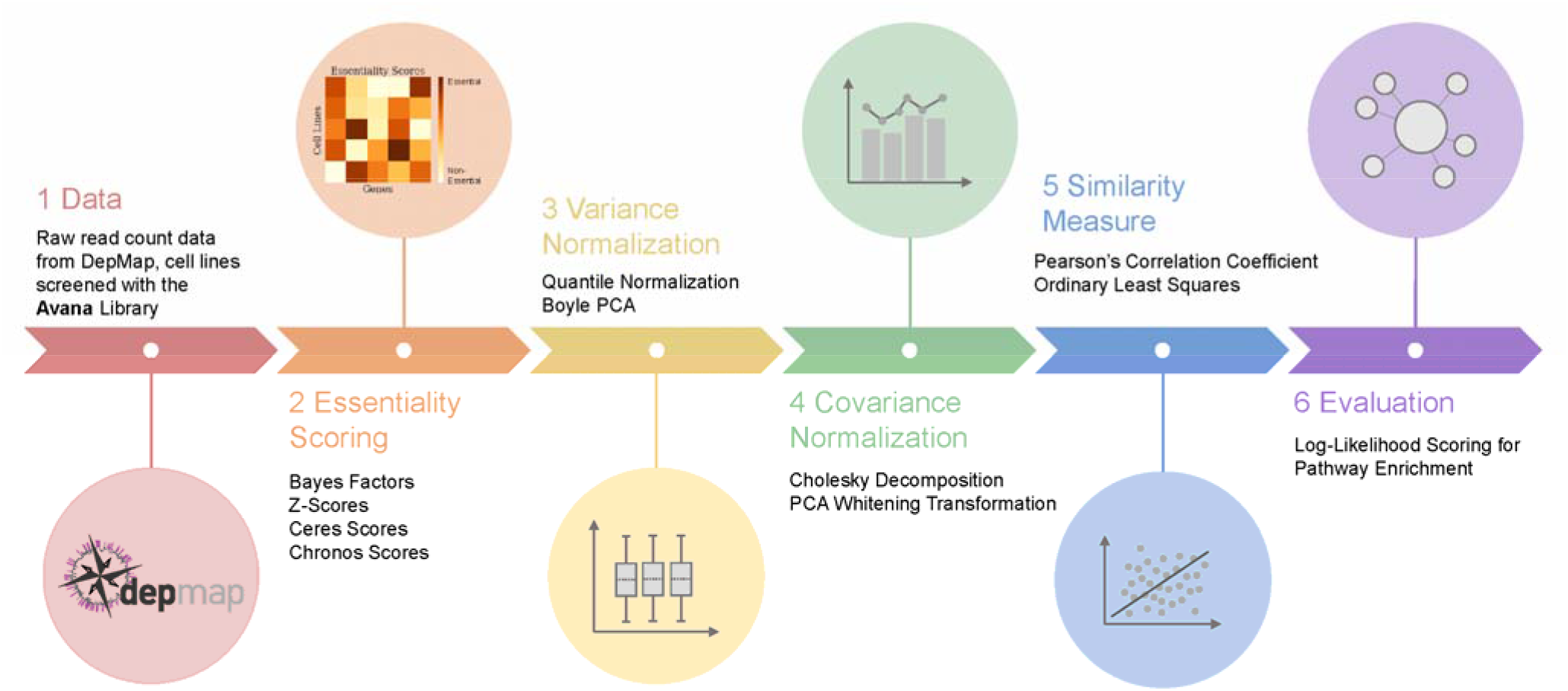
The network construction pipeline. We generated coessentiality networks using all combinations of essentiality scoring, variance normalization, covariance normalization, and similarity measurement methods, and each network is evaluated for pathway enrichment. All networks are built from the same raw read count data.

Gene knockout fitness effects (“essentiality scores”) were calculated using four recent algorithms designed explicitly for the analysis of CRISPR pooled library knockout screens. Bayes Factors (BF) were calculated using the BAGEL2 algorithm (Kim and Hart, 2021); Ceres (Meyers et al., 2017) and Chronos (Dempster et al., 2021) scores were downloaded from the Dep Map portal (Broad Institute, 2019), and Z-scores were calculated from a two-component Gaussian mixture model of average fold changes as described in Lenoir et al (Lenoir et al., 2021). We selected the intersection of all genes and samples measured by all algorithms, resulting in a matrix of 17834 genes by 730 cell lines, ensuring a comparable result from each algorithm.

We previously used quantile normalization as a method of sample-level variance normalization, where each gene’s fitness score is normalized to the rank mean (see Methods). An alternative approach described in Boyle et al (Boyle et al., 2018) that uses singular value decomposition of a reference set of nonessential genes to identify and remove “artifactual” components is functionally equivalent to sample-level variance normalization (Supplementary Figure 1) and is therefore included here.

Covariance normalization or “whitening” can be an important step to prevent correlated samples from biasing results; e.g. when specific tissue types or oncogenic mutations are overrepresented in the cell lines. PCA-based whitening is one such approach. Wainberg et al (Wainberg et al., 2021) introduce Cholesky decomposition as a covariance normalization in their coessentiality network which, coupled with ordinary least squares (OLS) regression, they apply as a generalized least squares (GLS) approach. We apply Cholesky whitening (covariance normalization) and OLS (similarity measure) separately to evaluate the relative contributions of each, and include the commonly used Pearson correlation coefficient (PCC) as an alternative similarity measure. See Figure 1 for an overview of the processing steps applied, and Methods for implementation details.

To evaluate the ability of a coessentiality network to identify co-functional gene pairs, we compared the most similar (top 50k) gene pairs from each network to annotated pathway databases. We measured enrichment by applying the log likelihood framework developed for functional interaction networks (Lee et al., 2004), where gene pairs belonging to the same annotated pathway are considered true positives and pairs belonging to different pathways are false positives (Figure 2A). We explored several pathway annotation databases, including Reactome (Croft et al., 2011), KEGG (Kanehisa and Goto, 2000), and the GO Biological Process tree (Ashburner et al., 2000), and determined that Reactome offered the most complete coverage (Supplementary Figure 2). Interestingly, we found that a very large number of genes and gene pairs in the annotated set belonged to just six annotated pathways involving mitochondrial translation and oxidative phosphorylation. Since the number of edges in a fully connected network grows with the square of the number of nodes, the large size of the mitochondrial ribosome and ETC Complex I result in a very large number of “true positive” hits that can overestimate the quality of the remaining network (Supplementary Figure 2). Since oxphos genes are known to be a source of bias in coessentiality networks and differential essentiality, we removed these six pathways from the Reactome reference set, hereafter referred to as CleanReactome.

**Figure 2.**
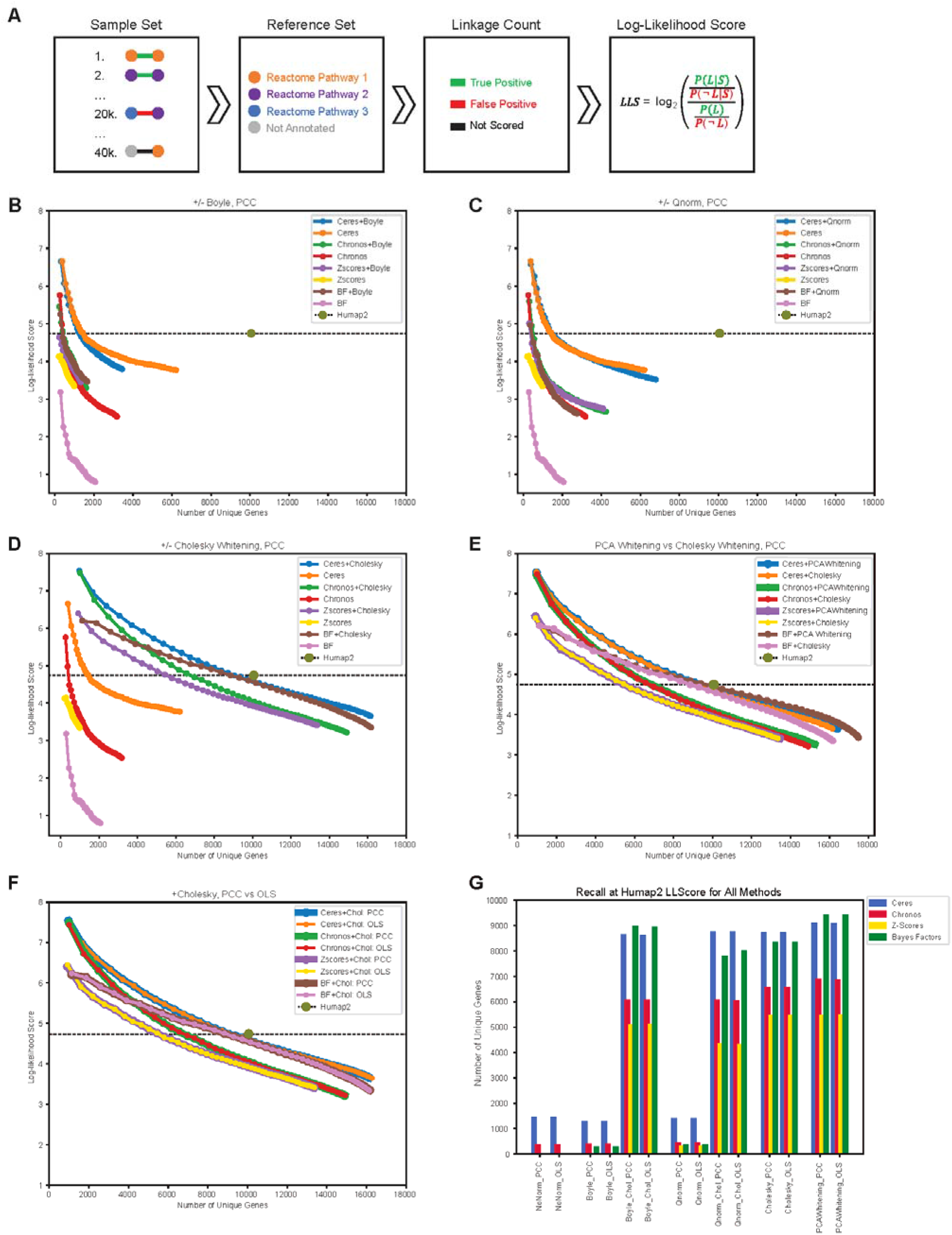
Evaluation of network construction methods. **A)** The Log-likelihood framework for evaluating a network for pathway enrichment. Gene pairs in the sample are compared against reference set CleanReactome. Pairs annotated in the same Reactome pathway are counted are true positives, while pairs annotated in different pathways are false positives. **B)** Top ranking 50k gene pairs from networks created with and without applying Boyle PCA normalization are binned in bins of 1000 pairs, and evaluated with LLS scheme. For each network, LL scores are plotted vs the number of unique genes in each bin, calculated cumulatively. **C)** LLS and recall for networks created with and without Quantile-normalization applied. **D)** LLS and recall for networks created with and without Cholesky Whitening. **E)** LLS and recall for networks where PCA-whitening was used as covariance normalizations, compared to networks where Cholesky-whitening was used. **F)** LLS and recall, for networks with Cholesky-whitening applied, using PCC as similarity measure vs using OLS. **G)** Recall for all 56 networks at the bin where the networks LLScore is approximately equal to the Humap2 LLS.

We evaluated the cumulative (Figure 2B-F) and local (Supplementary Figure 3) functional enrichment LLS for the top 50,000 gene pairs in each network, in bins of 1,000 gene pairs, and plotted coverage (total number of genes in the network) vs functional enrichment (LLS score). Both Boyle (Figure 2B) and quantile normalization (Figure 2C) significantly improved both the coverage and accuracy of the Bayes Factor and Z-score networks, but had lesser impact on the Ceres and Chronos data, likely because both of these methods contain a functional equivalent to sample variance normalization.

In contrast to sample variance normalization, global covariance normalization dramatically improved both the coverage and accuracy of networks derived from all four essentiality scores. Cholesky whitening (Figure 2D) increased recall and functional enrichment for both Ceres and BF networks to nearly match that of the hu.MAP 2.0 compendium of human protein complexes, derived from integration of numerous large-scale affinity purification/mass spectrometry and other protein-protein interaction data (Drew et al., 2021). PCA whitening (Figure 2E) closely matched the improvement of the Cholesky approach, with trivial incremental improvement in some cases.

A key conclusion of the Wainberg et al study is that GLS, implemented there as Cholesky whitening plus OLS, is superior to PCC-derived functional interaction networks. However, after covariance normalization, OLS is mathematically identical to Pearson correlation (see Methods). This is reflected in the identical LLS curves generated by PCC or OLS derived similarity scores after Cholesky whitening (Figure 2F).

We summarized each network’s performance by calculating its recall at the functional enrichment level offered by hu.MAP2 (Figure 2G). Raw or sample-normalized networks offered weak performance compared to pipelines that include covariance normalization. Unsurprisingly, sample variance normalization steps (Boyle, quantile normalization) provide no incremental improvement when covariance normalization is applied, OLS and PCC are identical after covariance normalization, and different covariance normalization methods give very similar results. Perhaps more surprising is the significantly better coverage of networks generated with Ceres or BF scores relative to Z-scores or the newer Chronos scores, the current method of choice for reporting DepMap hits.

Though these networks may yield equivalent scores of global accuracy, it is not necessarily the case that they contain identical information. To assess systematic differences between high-scoring networks, we examined differences between the covariance normalized Ceres network (top 17,000 edges) and the normalized BF (top 15,000 edges) and Chronos (top 12,000 edges) networks (Figure 3A). The Ceres and BF networks share only ~7,000 edges and ~6,500 genes, while each network contains more than 2,500 unique genes (Figure 3B). Common edges are generally correlated but both networks contain high-scoring edges that are unique to that processing pipeline (Figure 3C). However, the genes unique to each pipeline show no functional enrichment for GO or KEGG pathways, suggesting no functional bias. Likewise, the Ceres and Chronos networks share nearly 8,000 edges and 6,000 genes (Figure 3D) and common edges are highly correlated (Figure 3E). The Ceres network, larger because of its greater recall at the hu.MAP2 LLS threshold, has 3,202 unique genes that are highly enriched for core cellular processes such as transcription and translation, suggesting that these genes are depleted in the Chronos network (Figure 3F). This is not solely a result of the smaller Chronos network; a similar analysis of identical-sized networks yields the same results (Supplementary Figure 4), suggesting a systematic limitation of the Chronos scoring scheme relative to Ceres.

**Figure 3.**
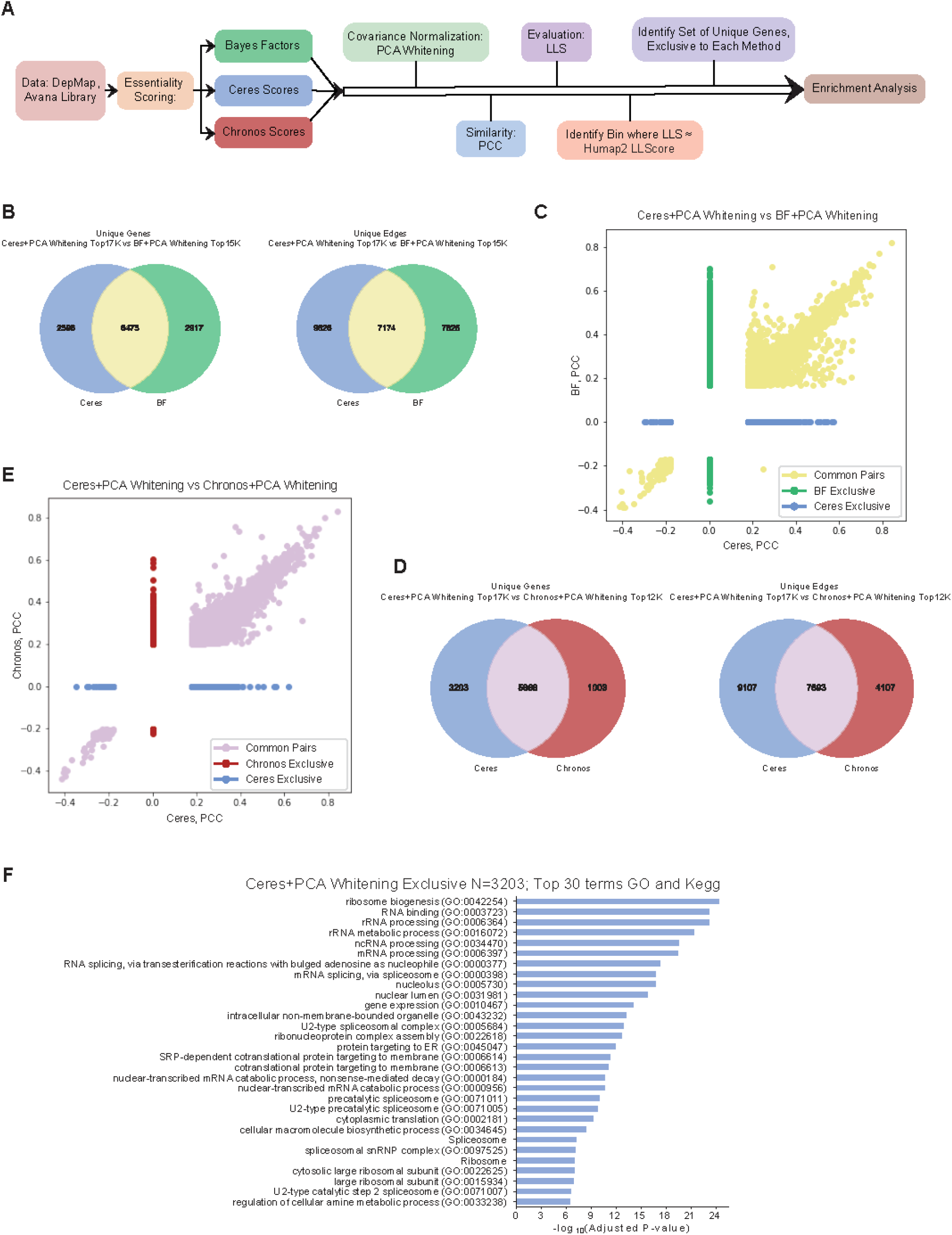
Comparison of top networks. **A)** Schema for enrichment comparison of top scoring networks. Ceres, BF, and Chronos scroes were subjected to identical processing pipelines and the resulting networks compared. **B)** Venn diagrams depicting the numbers of genes and edges exclusive to the top 17k edges in the network created with Ceres+PCAwhitening+PCC vs the top 15k edges in the network created with BF+PCAwhitening+PCC. **C)** Pearson’s correlation coefficients of common edges, and of exclusive edges to each of the networks (Ceres vs BF). **D)** Venn diagrams depicting the numbers of genes and edges exclusive to the top 17k edges in the network created with Ceres+PCAwhitening+PCC vs the top 12k edges in the network created with Chronos+PCAwhitening+PCC. **E)** Pearson’s correlation coefficients of common edges, and of exclusive edges to each of the networks (Ceres vs Chronos). **F)** Enrichment of the gene set exclusive to Ceres+PCAwhitening+PCC (when compared to Chronos). Only the top 30 enriched GO and Kegg terms are listed in the graph.

Wainberg et al argue that the GLS approach, combining covariance normalization with OLS, is superior to PCC because the P-values of PCC networks can be inflated by unequally weighted samples (Wainberg et al., 2021), leading to false positives. However, PCC and OLS are mathematically equivalent after covariance normalization, and this is reflected in the identical performance curves of the PCC and OLS similarity measures as shown in Figure 2F. One advantage of Pearson correlation is that it is far less computationally intensive; an all-by-all gene correlation matrix takes only a few seconds (and one line of code) to calculate while OLS requires setting up the regression for each gene pair individually, which can take orders of magnitude longer. Notably, global PCC values after whitening are significantly smaller in magnitude than PCC before whitening, leading to some counterintuitive results; at the same rank, post-whitening PCC is often considerably weaker than pre-whitening PCC (Supplementary Figure 5) despite showing a significantly higher enrichment for functional interaction (Figure 2D).

A second advantage of the Pearson correlation approach is that it facilitates the use of partial correlation to detect conditional interactions between sets of genes. Partial correlation can measure the similarity of two gene knockout profiles after removing the effect of a third gene, set of genes, or other vector. If the functional relationship between two genes varies across the assayed cell lines, the observed correlation can be diluted by the presence of samples in which the relationship is severed (Kim et al., 2022b). For example, consider common oncogenes *KRAS* and *NRAS*, two members of the MAP kinase signaling pathway whose mutually exclusive mutations mean they are essential in different cell lines (Figure 4A). Downstream signaling partner RAF1 is also essential in the presence of either *KRAS* or *NRAS* (Figure 4A). The PCC of *RAF1-NRAS ρ_RAF1,NRAS_* is significant, ranking in the top 0.003% of global correlation pairs, but the partial correlation of RAF1-NRAS with respect to KRAS *ρ_RAF1,NRAS_KRAS_* is even higher. Likewise, *ρ_RAF1,KRAS_NRAS_* > *ρ_RAFl,KRAS_*, indicating higher correlation after removal of the NRAS signal, while the mutual exclusivity of KRAS and NRAS mutations results in *ρ_KRAS,NRAS_* < 0 (Figure 4B). This can be considered a case of a protein “moonlighting” where *RAF1* interacts with either *KRAS* or *NRAS* depending on the context.

**Figure 4.**
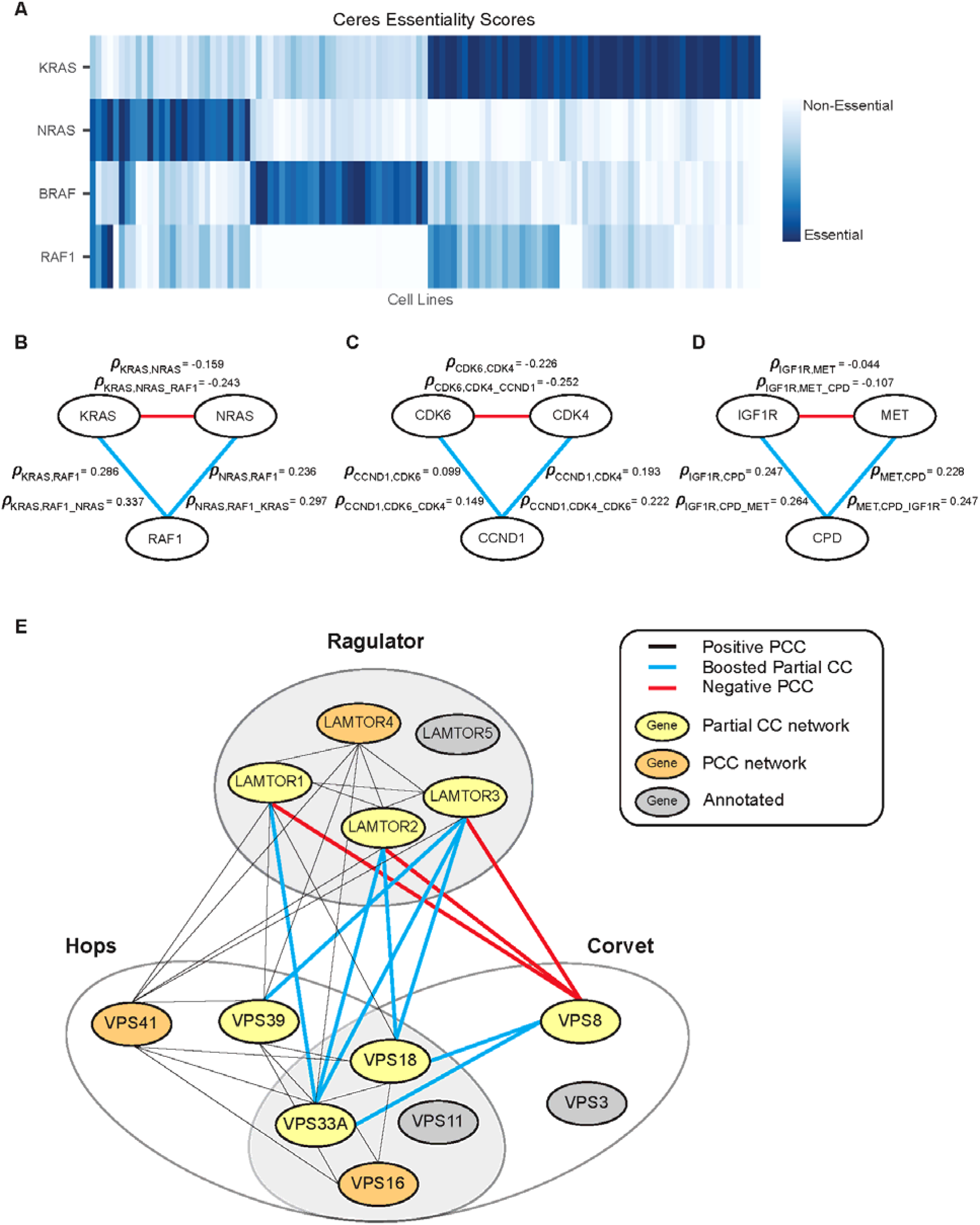
Detecting moonlighting interactions using partial correlation. **A)** Heatmap showing KRAS, NRAS, BRAF essentiality in selected cell lines. KRAS, NRAS, and BRAF are mutually exclusive, while RAF1 is essential in KRAS and NRAS backgrounds. **B)** KRAS-NRAS-RAF1 moonlighting trio, with PCC and Partial correlation coefficients listed by the respective edges. Red edges indicate negative correlation coefficient, while blue edges indicate positive correlation coefficient. **C)** CDK6-CDK4-CCND1 moonlighting trio, with correlation coefficients of edges listed. **D)** IGF1R-MET-CPD moonlighting trio, with correlation coefficients of edges listed. **E)** Network illustrating Ragulator complex, HOPS and CORVET complexes with shared subunits. Red colored edges indicate negative PCC, blue colored edges indicate boosted partial correlation coefficient, black colored edges are positive PCC. The yellow nodes depict genes recovered using partial correlation, orange nodes are genes recovered with PCC, and gray nodes depict genes annotated to be in the complex, that were not present in our top-ranking pairs.

We conducted a systematic search for these moonlighting trios, where the candidate moonlighting gene has a positive PCC with each candidate “parental” gene, the partial correlation with each parent gene is boosted after controlling for the other parent, and the PCC of the two parents is negative. We discovered several cases that reflect known examples of context-dependent interaction. D cyclin *CCND1* interacts with cyclin-dependent kinases *CDK4* and *CDK6*, which are mutually exclusive in DepMap data (Figure 4C). Similarly, protease *CPD* interacts independently with insulin-like growth factor receptor *IGF1R* and hepatocyte growth factor receptor *MET* (Figure 4D). CPD protein is known to process pro-IGF1R into its mature form (Han et al., 2020) and we recently showed that *CPD* is also involved in maturation of *MET* receptor protein specifically in glioma cells (Kim et al., 2022).

More complicated conditional interaction subnetworks can also be informative, including at least one instance where partial correlation can disambiguate protein complexes with overlapping membership (Figure 4E). The CORVET and HOPS complexes are functionally related molecular machines that play important roles in endocytosis (Solinger and Spang, 2013), with the CORVET complex being primarily involved in early endocytosis and being replaced by HOPS at the late endosome/lysosome stage (Balderhaar and Ungermann, 2013). The Ragulator complex is a multisubunit complex that sits at the lysosome and acts as an activator of mTORC1 complex in the presence of amino acids, and is itself regulated by the HOPS complex (Sancak et al., 2010). Partial correlation analysis differentiates shared CORVET/HOPS subunits VPS33A and VPS18 from CORVET-specific subunit VPS8, while their partial correlation with Ragulator subunits LAMTOR1-4 are boosted after removing VPS8 effects (Figure 4E). HOPS-specific subunit VPS39 is correlated with the Ragulator independent of CORVET-specific VPS8.

## Conclusions

Coessentiality networks offer a powerful method for inferring gene function from panels of CRISPR knockout screens in cell lines, but “coessentiality” is a catchall term describing an informatic pipeline with a variety of choices. We systematically explored the most common of these options to determine which combination of fitness scores, variance normalizations, and similarity measures gave maximal coverage of known co-functional relationships. We found that covariance normalization or “whitening” gives the largest boost to performance, regardless of fitness scoring, but also that coessentiality networks derived from covariance-normalized Bayes Factor and Ceres fitness scores markedly outperformed both the Chronos and Z-score approaches. Interestingly, after covariance normalization, the top two similarity measures, Pearson correlation and Ordinary Least Squares (OLS), are mathematically identical and give the same ranking for gene pairs.

Though genes which operate in the same biological process or pathway have similar knockout fitness profiles, genes whose functional interaction is context-dependent can have their profile similarities weakened by inclusion of cell lines or contexts where the interaction is not present. We (Kim et al., 2022) and others (Pan et al., 2022) have recently shown how analysis of CRISPR genetic screen data can reveal proteins that have multiple roles in a cell, leading to functional interactions with mutually exclusive partners. We extend the Pearson correlation network by using partial correlation to identify these “moonlighting” genes, whose interactions with one gene are boosted when the effect of a second gene is removed (and vice versa).

This work reinforces studies that show that coessentiality networks are among the most powerful predictors of mammalian gene function. Properly constructed, these networks contain more than 10,000 unique genes -- more than half of the protein-coding genome -- despite individual screens rarely recording more than 2,000 genes with fitness defects and entire categories of genes, e.g. those involved in secretory pathways or cell-cell communication, being systematically absent from pooled library screens. Moreover, the accuracy of these networks is comparable to that of the hu.MAP integrated map of protein complexes. Nevertheless, the systematic discovery and elucidation of context-dependent and pleiotropic gene functions across the DepMap cell lines has only just begun, and promises to increase our insight into the organization and function of mammalian cells.

## Methods

### Data and Essentiality Scoring

The data used in this study comes from publicly available CRISPR knockout screens datasets, downloaded from the Cancer Dependency Map database (Avana dataset) (Broad Institute, 2019). Four different pipelines for measuring gene knockout fitness effects (gene essentiality) were used: BAGEL2 (Kim and Hart, 2021), Z-score model (Lenoir et al., 2021), Ceres (Meyers et al., 2017), and Chronos (Dempster et al., 2021). The BAGEL2 algorithm generates log Bayes Factors (BF) to report gene essentiality for each cell line, with positive scores indicating essentiality. Z-score values are generated using a Gaussian mixture model, with negative scores indicating essentiality. The Ceres algorithm removes principle components highly related to copy-number-specific effects and scales the data so that median essential score is −1, and median non-essential score is 0. Chronos models the read-count data assuming a negative binomial distribution and removes copy-number related bias, with the scores scaled similar to Ceres.

Bayes Factors (BF) and Z-scores were calculated using raw read count data from the DepMap 20Q4 release. Ceres scores were downloaded from the 20Q4v2 release, and Chronos gene effect scores were downloaded from the 22Q2 release. We considered only the common genes and cell lines of BF, Z-scores and Ceres scores, which resulted in genes-by-cell lines data matrices of size 17834 × 730. The Chronos data was processed to include only genes and cell lines present in the intersection with Ceres, BF and Z-score, resulting in a data matrix of size 17662 × 727.

### Normalization Techniques

We compared four normalization techniques, two of which perform as variance normalization methods and the other two are covariance normalization methods. The quantile normalization technique executes variance normalization, applied to mitigate screen quality bias and to allow comparison between different samples. Quantile normalization first ranks the genes by magnitude, calculating the mean for genes in the same rank, and substituting the values of all genes in that rank with the mean value, and then reorders the genes in the original order. The Boyle principal component analysis approach aims to remove the technical confounding introduced by olfactory receptor genes that have highly correlated profiles across different genetic backgrounds (Boyle et al., 2018). This method applies principal component analysis of the gene-by-cell-line essentiality matrix across olfactory receptors, and then subtracts the first four principal components from the original score matrix.

The covariance normalization methods, also known as whitening or sphering transformations, aim to remove dependencies between features in the data matrix. These methods involve linear transformation of the data matrix *X* using a “whitening” matrix *W*, such that the resulting normalized data matrix 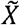 has a covariance matrix equal to the identity matrix (Eqn 1). To perform this, the data matrix is first centered, subtracting the mean across all samples, and the covariance matrix is computed, denoted by Σ (Eqn 2). The covariance matrix is positive semi-definite, meaning it is symmetric with non-negative eigenvalues, thus it can be inverted, and it can be decomposed as a product of two simpler matrices. There are many “whitening” matrices that can do the linear transformation described above, and we used two different options that satisfy the condition. One of the techniques of calculating *W*, described in Wainberg et al (Wainberg et al., 2021), uses a Cholesky decomposition of the inverse covariance matrix (Eqn 3), and the linear transformation is done as described in (Eqn 4). Another technique termed PCA whitening utilizes the Eigen-decomposition of the covariance matrix (Eqn 5), and the PCA whitening transformation is done as described in (Eqn 6).

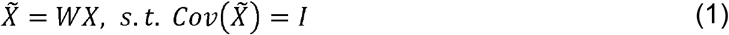

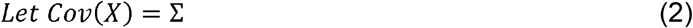

Cholesky Whitening:

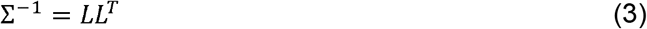

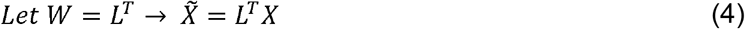

PCA Whitening:

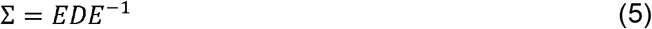

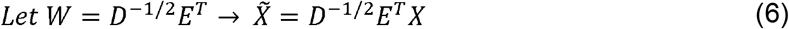

Where *D* = *diag*(*λ*), a diagonal matrix containing the eigenvalues *λ_i_* on the diagonal; and E is the orthogonal matrix of eigenvectors.

### Statistical Measures of Similarity

We utilized two statistical measures to quantify co-essentiality, Pearson’s correlation and Ordinary Least Squares. Using Pearson’s correlation, we calculate the correlation coefficient for all possible gene pairs (Eqn 7), resulting in a gene-by-gene correlation matrix. We rank the gene pairs by the correlation coefficient values.

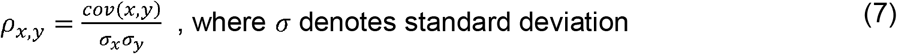

Using ordinary least squares (numpy.linalg.lstsq), we estimate the parameter vector *b* in the linear regression model *y* = *bx* + *ε*, where *y* is set to be the essentiality scores for one of the genes and *x* is a two-column matrix, with the first column being the other gene’s essentiality scores and the second column is the intercepts, set to a vector of all ones. We ran OLS for each gene pair, and calculated log P-values, resulting in a gene-by-gene matrix of P-values. When the Cholesky whitening transformation is applied, both *x* and *y* are transformed by the triangular Cholesky matrix, and this constitutes the Generalized least squares method described in Wainberg et al. We rank the gene pairs by the log P-values.

It is important to note that in least squares linear regression the slope *b* is given by (Eqn 8). Rewriting it as in (Eqn 9) and substituting (Eqn 7), shows how the correlation coefficient *ρ* factors in. When covariance normalization is applied, whether with Cholesky or PCA whitening, the transformed data has identity covariance, meaning all features are independent and the variance along each of the features is one. With the variance (*σ*^2^) equal to one in the covariance normalized data, PCC and OLS yield equivalent results.

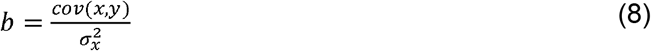

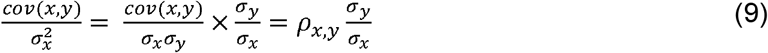

### Evaluation of Pathway Enrichment with Log-Likelihood Scoring

For each method, we took the top 50,000 ranking gene pairs, and bin them into bins of 1,000 gene pairs for further analysis. To evaluate the performance of these methods, we conducted pathway enrichment tests with a log-likelihood scheme described by Lee et al (Lee et al., 2004), using multiple annotated pathway databases. The pathway databases we considered are: Kegg (Kanehisa and Goto, 2000), Gene Ontology (GO) Biological Processes (Ashburner et al., 2000), and Reactome (Croft et al., 2011). Additionally, we considered pre-processed versions of GO and Reactome, in which pathways that contained gene sets involved in mitochondrial translation and oxidative phosphorylation were removed. We refer to these versions as CleanGO and CleanReactome. Only six pathways were removed from Reactome in this process, and the bulk of our analysis was done with the resulting CleanReactome. In the log-likelihood scoring pipeline, we filter through the pathways within a reference file, to only include those pathways that have a minimum of 5 genes and maximum of 400 genes.

Using the gene pairs from each bin and the annotated reference set, we measure if both genes in the sample pair are annotated in the same pathway (true positive), or if they are annotated in different pathways (false positive). If a gene from our sample is not annotated in any pathways in the reference set, we do not count the pair in our calculation. We then compare the ratio of the frequencies of true positives and false positives in our sample with the background expectation, meaning the ratio of frequencies of all annotated genes operating in the same pathway and genes operating in different pathways in the reference set. The log-likelihood score is given by (10).

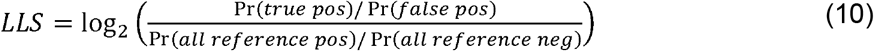

Scores greater than zero indicate that the method tends to link genes in the same pathway, with higher scores indicating more confident linkages. The log-likelihood scoring was performed cumulatively, as well as locally for each method.

The hu.MAP 2.0 Protein Complexes List was downloaded from the humap2 website (http://humap2.proteincomplexes.org/static/downloads/humap2/humap2_complexes_20200809.txt) (Drew et al., 2021). We created a list of unique gene pairs from each protein complex, a total of 57,911 gene pairs, and calculated overall LLS for this list. The LL score for hu.MAP 2.0 was 4.75, with coverage of 10,060 unique genes. We used this score as a reference against which to compare the performance of the networks formed with the various methods.

### Gene Set Enrichment Analysis

To analyze the differences between the highest scoring networks, we looked at the enrichment from genes exclusive those networks. We used the hu.MAP2 LL Score as a cut-off for the PCA-whitening covariance normalized Ceres, Bayes Factors, and Chronos networks. We identified edges and genes exclusive to each network, and conducted enrichment analysis on the exclusive gene sets using the GSEAPY “enrichr” python module (Kuleshov et al., 2016) with the reference sets ‘KEGG_2021_Human’, ‘GO_Biological_Process_2021’, ‘GO_Cellular_Component_2021’, ‘GO_Molecular_Function_2021’.

### Partial Correlation

We utilized partial correlation to reveal the conditional relationship between two genes, after controlling the effect of a third gene. We calculate partial correlation of two genes *x* and *y*, while controlling the effect of a third gene *z*, using the recursive formula (Eqn 11). For each pair of genes *x* and *y* with |*ρ_x,y_*| > 0.15, we calculate partial correlation with respect to every other gene in the network, and look for the gene that yields the highest partial correlation coefficient. To quantify the change in correlation coefficient after accounting for the effect of a third gene,

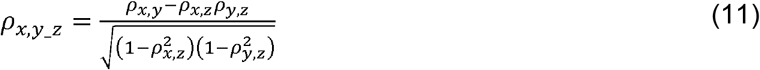

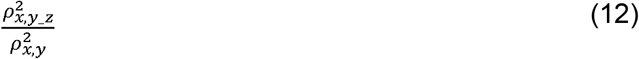

we calculate a ratio between the two coefficients with the formula (Eqn 12).

Our partial correlation analysis revealed many gene trios (*x-y-z*), where positive correlation coefficients between genes *x* and *y* and between genes y and z are boosted after controlling for the effect of the other, while the correlation coefficient between genes *x* and *z* is negative. A master data list of these trios is available in Supplementary Table 1. We created a network with these moonlighting trios, consisting of gene pairs with the edge being the positive partial correlation coefficient, as well as gene pairs with negative PCC edge. The network can be viewed in Cytoscape (Shannon et al., 2003), and the data table used to create the network is available as a Supplementary Data file.

## Acknowledgments

VG and TH were supported by NIGMS grant R35GM130119. TH is a CPRIT Scholar in Cancer Research (RR160032). This work was supported by the NCI Cancer Center Support Grant P30CA16672.

## Author Contributions

VG developed the approach and performed all bioinformatic analysis. VG and TH drafted and edited the manuscript.

