## Supplementary Figures for "Optimal construction of a functional interaction network from pooled library CRISPR fitness screens"

Supplementary Figure 1.

**A**

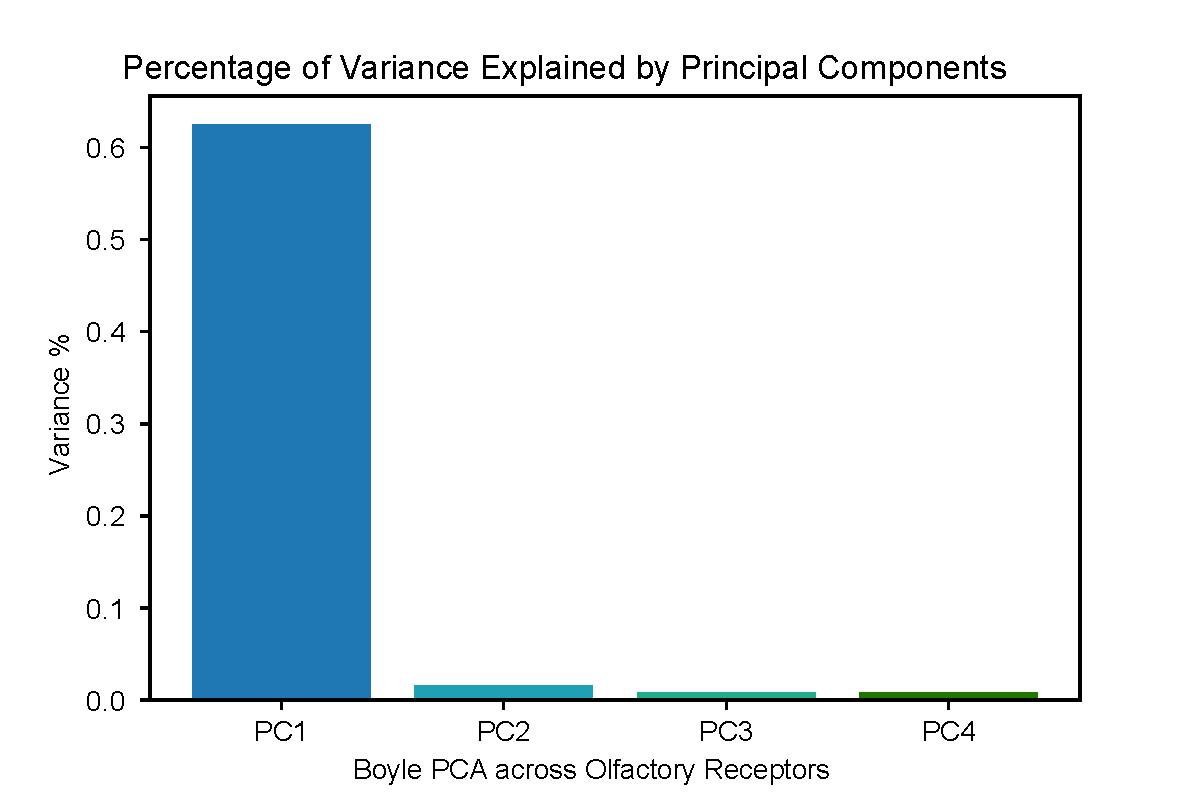

**B**

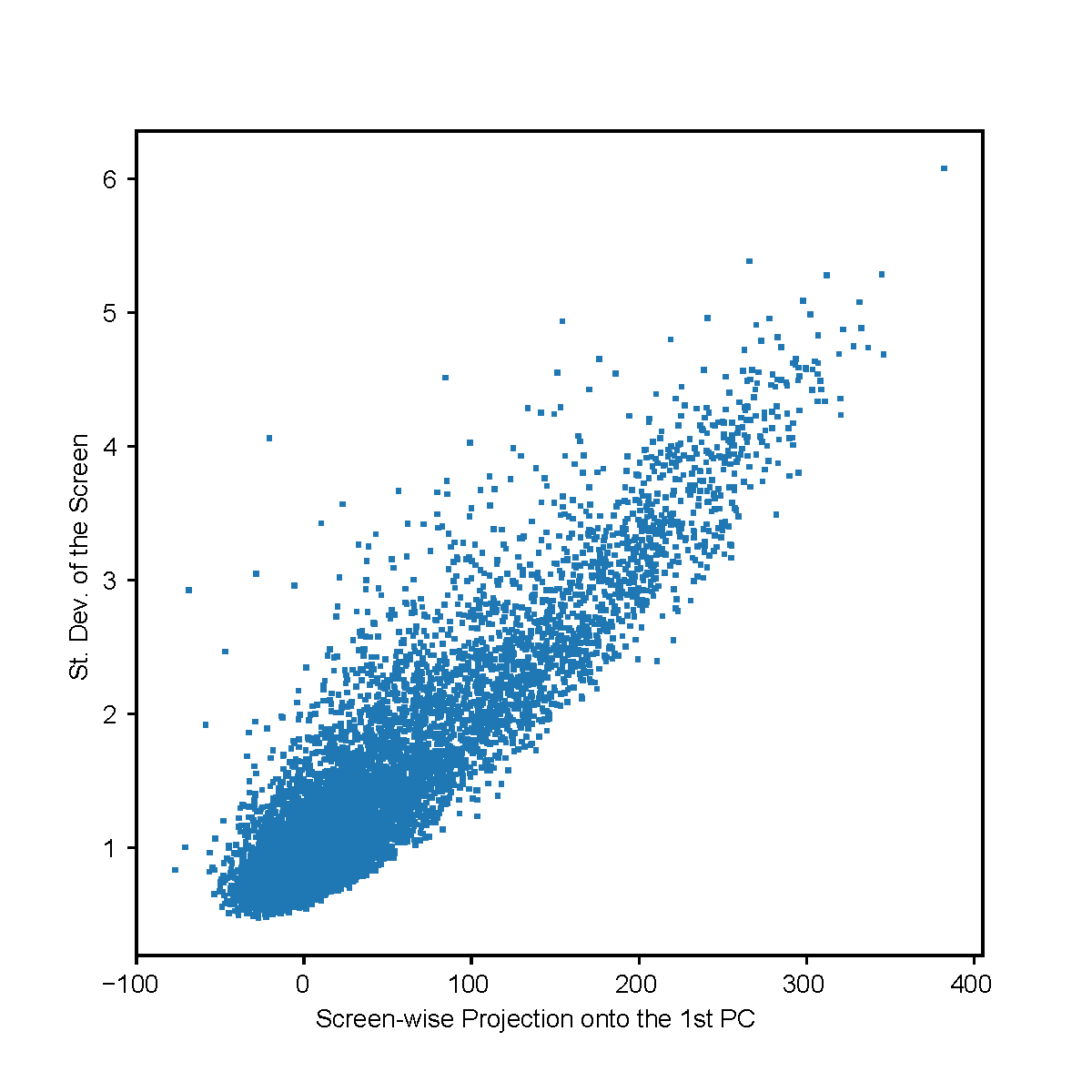

**Supplementary Figure 1.** **A)** Bar plot showing the percentage of variance explained by each principal component of the Boyle PCA approach across Olfactory receptor genes, applied to the Z-score data matrix. **B)** Scatter plot of the standard deviation of the screen, using Z-scores data matrix, versus the screen-wise projection onto the first Principal component from the Boyle approach.

Supplementary Figure 2.

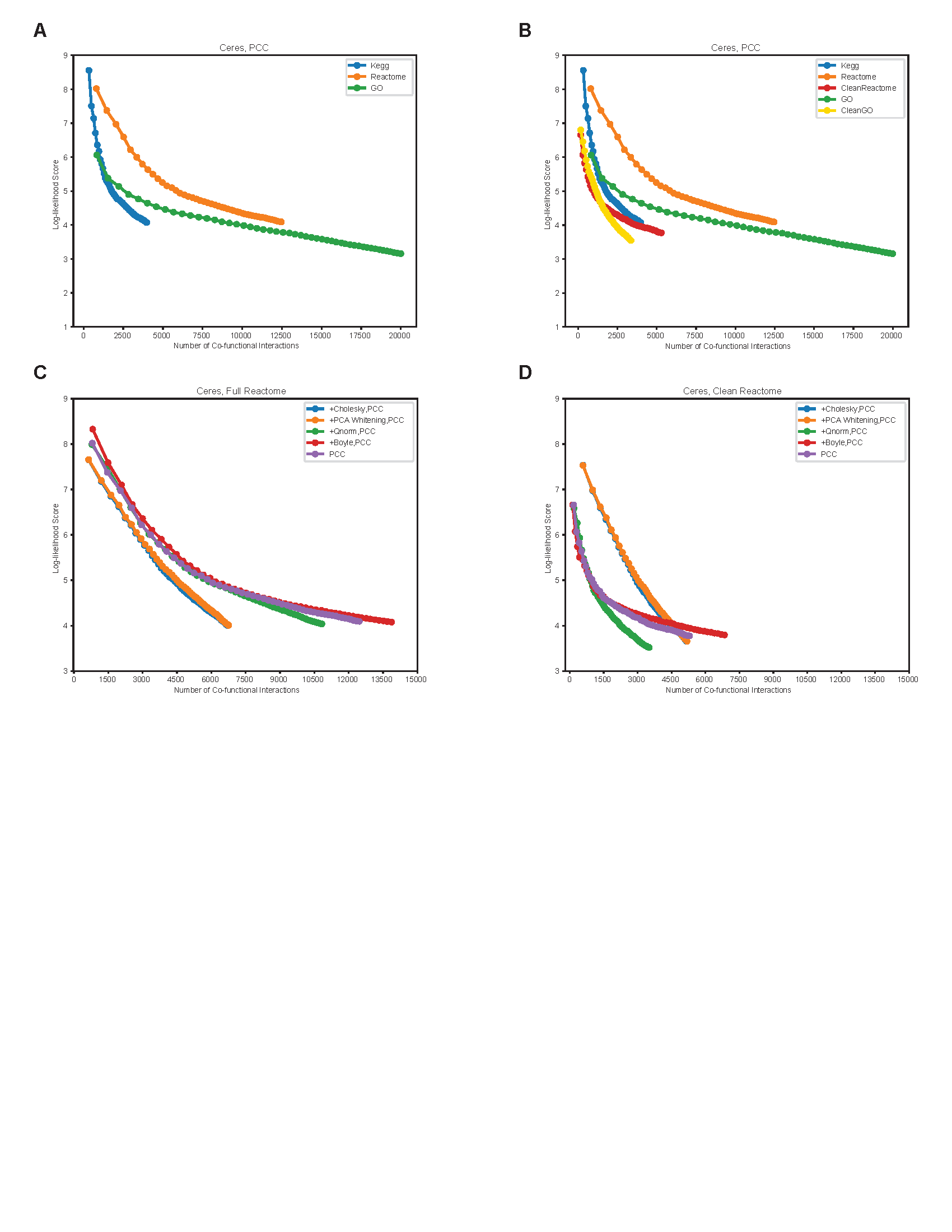

**Supplementary Figure 2.** **A)** Ceres+PCC network evaluated using 3 different reference sets in the Log-likelihood analysis. Cumulative LL Scores and number of co-functional interactions in bins are plotted for the network, evaluated with Kegg, Reactome and GO reference sets. **B)** Pathways containing genes associated with mitochondrial translation and oxidative phosphorylation were removed from Reactome and GO, to create CleanReactome and CleanGO. Cumulative LL Scores and number of co-functional interactions in bins are plotted for the Ceres+PCC network, evaluated with Kegg, Reactome and CleanReactome, GO and CleanGO reference sets. **C)** Comparison of LLS scores and co-functional interactions of Ceres based networks using the full Reactome reference set and **D)** the CleanReactome reference set in the LLS evaluation.

Supplementary Figure 3.
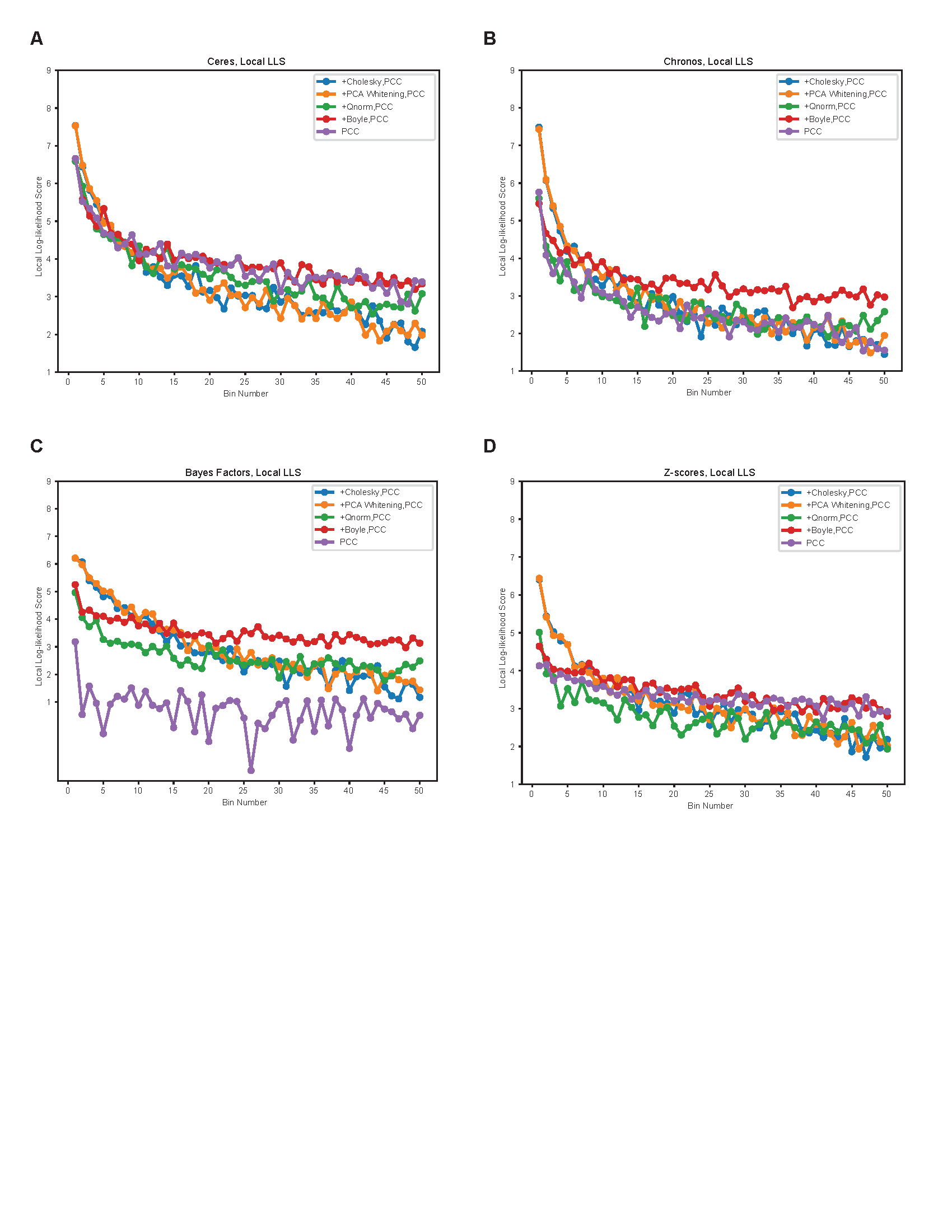

**Supplementary Figure 3**. **A)** Local log-likelihood scores calculated per bin, using CleanReactome, for Ceres-based networks; **B)** Chronos-based networks; **C)** Bayes Factors based networks and **D)** Z-scores based networks.

Supplementary Figure 4.

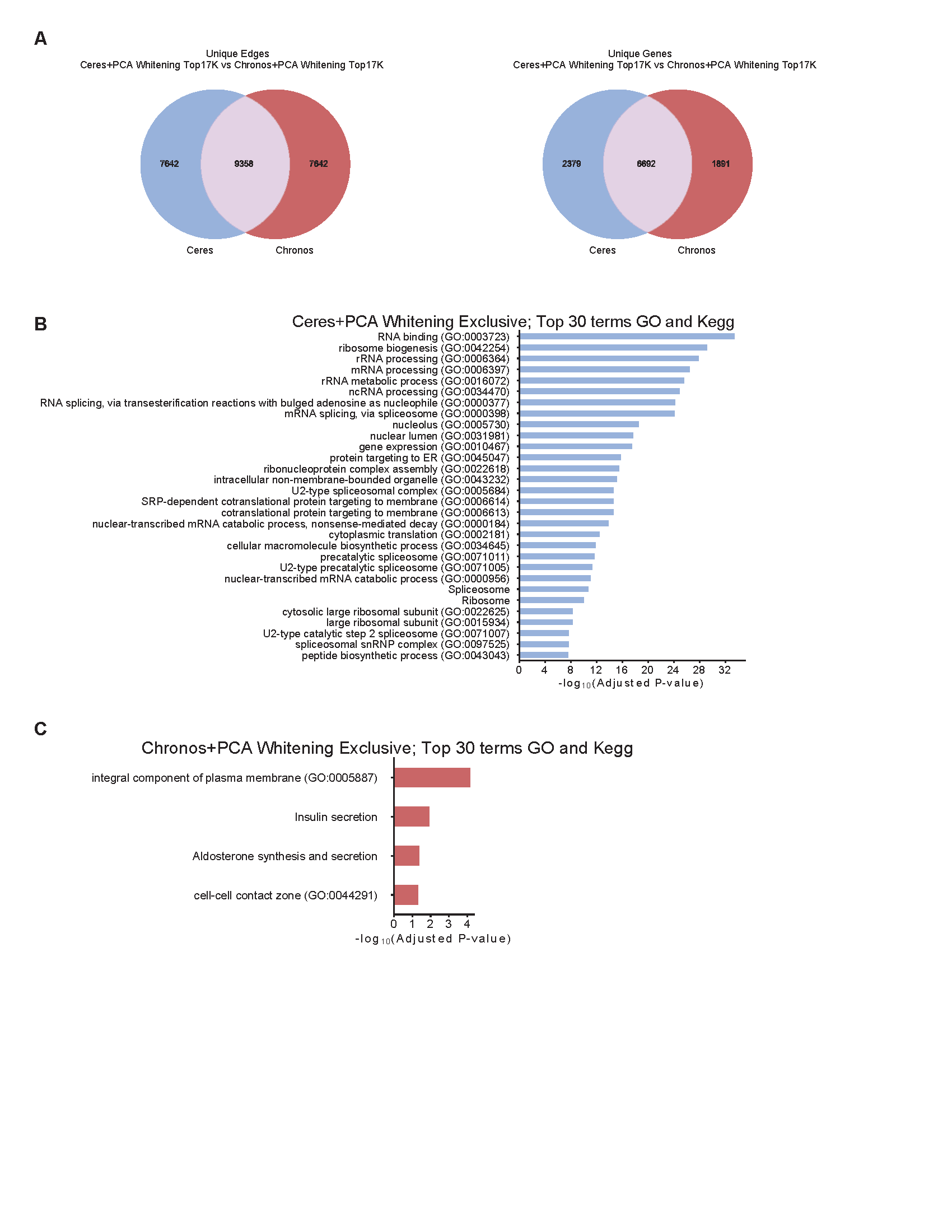

**Supplementary Figure 4**. **A)** Venn diagrams depicting the numbers of genes and edges exclusive to the top 17k edges in the network created with Ceres+PCAwhitening+PCC vs. the top 17k edges the network created with Chronos+PCAwhitening+PCC. **B)** Enrichment of the gene set exclusive to Ceres+PCAwhitening+PCC. Only the top 30 enriched GO and Kegg terms are listed in the graph. **C)** Enrichment of the gene set exclusive to Chronos+PCAwhitening+PCC.

Supplementary Figure 5.

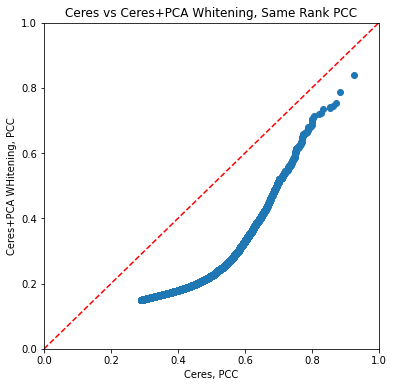

**Supplementary Figure 5**. Pearson’s correlation coefficients of the edges of the same rank in the Ceres+PCAwhitening+PCC and Ceres+PCC networks.
